# Individual differences in social reward and threat expectancies linked to grey matter volumes in key regions of the social brain

**DOI:** 10.1101/2020.02.03.916999

**Authors:** Bonni Crawford, Nils Muhlert, Geoff MacDonald, Andrew D. Lawrence

## Abstract

Prospection (mentally simulating future events) generates emotionally charged mental images that guide social decision-making. Positive and negative social expectancies – imagining new social interactions to be rewarding vs. threatening – are core components of social approach and avoidance motivation, respectively. Stable individual differences in such positive and negative future-related cognitions may be underpinned by distinct neuroanatomical substrates. Here, we asked 100 healthy adults to vividly imagine themselves in a novel self-relevant social scenario that was ambiguous with regards to possible social acceptance or rejection. During this task we measured their expectancies for social reward (e.g. anticipated feelings of social connection) or threat (e.g. anticipated feelings of rejection). On a separate day they underwent structural MRI; voxel-based morphometry (VBM) was used to explore the relation between their social reward and threat expectancies and regional grey matter volumes (rGMV). Increased rGMV in key regions involved in prospection, subjective valuation and emotion regulation (including ventromedial prefrontal cortex), correlated with both higher social reward and lower social threat expectancies. In contrast, social threat expectancies were uniquely linked with rGMV of regions involved in social attention (posterior superior temporal sulcus) and interoception (somatosensory cortex). These findings provide novel insight into the neurobiology of future-oriented cognitive-affective processes critical to adaptive social functioning.

## Introduction

Making friends - and/or forming romantic partnerships – is of critical importance for adults’ adjustment to new environments, for instance, starting university.^1^ Friendship bonds are consistently shown to have equal, or even greater, importance than family ties in predicting psychological well-being and physical health in adulthood.^1–8^ Research that has looked at not just the quantity, but also the quality, of social bonds has demonstrated that the mere existence of social relationships does not necessarily contribute positively to health.^4,9–11^ Supportive and rewarding social connections exert powerful effects on health and wellbeing, but relationship strain and social distress – the extent to which an individual perceives their daily social interactions as negative or distressing – can have equally strong, deleterious effects on health and wellbeing.^12,13^ Thus, social interactions and relationships are strongly linked to health and well-being because they present the potential for powerful (emotional) rewards as well as the potential for potent threats.^14,15^

Chosen relationships typically emerge from interactions among people. Humans are therefore intrinsically motivated to actively seek out and affiliate with others, with the aim of fostering new social connections.^16^ By their nature, however, social interactions with unfamiliar others simultaneously offer the prospect of both rewards (e.g. having a pleasant conversation, feeling a sense of belonging)^17^ and threats (e.g., feeling embarrassed, being socially rejected).^18,19^

Frameworks describing general motivation posit two basic systems that mediate actions geared towards desirable and undesirable outcomes – an approach (or behavioural activation) system (BAS) and an avoidance (or behavioural inhibition) system (BIS), respectively.^20–22^ These are suggested to be independent, but jointly operating, neurobehavioral systems. Models of social motivation connect these basic approach/avoidance motivational processes with social cognition, including attentional focus and beliefs about other people’s behaviour in social interactions.^2,14,15,23,24^

General approach and avoidance motivation are stable dispositions, albeit with some variation in adolescence and young adulthood.^25–27^ Sensitivity to social reward and threat are similarly stable,^28^ although both appear to be heightened during adolescence.^29^ These stable traits are associated with the likelihood of being socially connected or, conversely, isolated.^3^ It has been suggested that sophisticated neural-cognitive systems for calibrating social approach/avoidance motivation (and behaviour) evolved as a means of regulating hierarchies in complex primate societies.^30^ It seems plausible that individual differences in these neurocognitive systems might exist on continua of shyness and sociability, respectively, with the extreme ends of these continua being clinically relevant.^31,32^ For instance, maladaptations of these systems could result in social anxiety (excessively high BIS^31^), social anhedonia (excessively low BAS^33^), or hypersociability.^34,35^ All of these conditions are associated with loneliness^32,36–40^ and with poorer health and wellbeing more generally.^37,38,41^

An emerging literature details neural responses to rejection or connection experiences and visual cues of social reward or threat.^42–45^ However, there is good reason to think that ***prospective*** cognitive-affect representations are at the heart of these putatively distinct social reward and threat motivational systems. BAS or BIS have been theorized to be primarily future-oriented (e.g. mediating hopes and fears about future desirable or undesirable outcomes^46–48^). Similarly, MacLeod^46,47^ argues that affect is directly related to cognition and that positive and negative future-related cognitions may best be perceived as two separate dimensions of experience. Such future-oriented emotion systems depend on the capacity for “mental time travel” inherent in episodic memory.^49,50^ Mental time travel enables vivid, detail-rich simulations of future events based on the flexible re-combination of episodic memories and newly generated images constructed by drawing on both episodic and semantic memory (e.g. beliefs, goals). Through the vivid imagination of future events, humans generate embodied predictions of events’ emotional impacts before their occurrence, which act as powerful motivators of goal-directed behaviour.^50,51^

While some recent research has studied individual differences in anticipated social reward and threat separately (e.g.^45,52,53^), to our knowledge, no neuroimaging research has directly examined *both* individual differences in future-oriented social reward and threat expectancies in the context of fostering new social connections. Building on work in the domain of close relationships^54–57^ we developed a new instrument to examine inter-individual differences in reward and threat expectancies in the context of a social interaction with unfamiliar peers. This novel measure, the levels of dispositional expectancies for social threat and reward scale (LODESTARS), asks participants to vividly imagine that they have joined a new group, club, or society, and that later that evening, they will be meeting other people in this group/club/society for the first time. Participants then make predictions about the probable emotional consequences of interactions and report their anticipatory and anticipated emotions, by responding to items such as “*I will probably meet one or more people who I will like a lot*”. The imagined scenario is ambiguous, in that it simultaneously holds the possibility for social reward and social threat, thus maximizing opportunities for individual differences to emerge.^58,59^

Individuals’ social reward and threat expectancies as measured by the LODESTARS are stable over time, are associated with other stable affective traits such as self-esteem, and may be grounded in temperament and attachment experiences.^60,61^ Given this trait-like stability, we predicted that individual differences in expectancies for social threat and reward would be associated with stable, structural aspects of the brain. Recent structural magnetic resonance imaging (sMRI) studies indicate that a number of social traits are reflected in brain macrostructure (regional grey matter volume, rGMV) as assessed by voxel-based morphometry (VBM).^62^ Here, we utilized VBM and an unbiased, whole brain analysis, to investigate the possibility of unique and overlapping rGMV correlates of inter-individual differences in social threat and reward expectancies (STE and SRE, respectively) as measured by the LODESTARS.

This was primarily an exploratory study. However, we made two tentative predictions, based on previous research. First, we predicted that ventromedial prefrontal cortex (vmPFC) would correlate positively with SRE and negatively with STE. vmPFC is involved in the construction of episodic memories and imagined future events, as well as their valuation based on current needs and goals.^63^ vmPFC activity scales with anticipated positive value.^64^ Given that specific functional tasks correlate with volumes of regions subserving those tasks, we predicted increased vmPFC volume linked to increased SRE.

Another well-established role of vmPFC is in the regulation of negative affect.^65,66^ Previous work has found that more successful emotion regulation is associated with greater rGMV in vmPFC,^67^ so we expected increased vmPFC volume to also relate to lower STE.

Secondly, we expected that rGMV of the amygdala and posterior superior temporal sulcus (pSTS) would be positively correlated with STE. An abundance of research implicates the amygdala in threat processing and, of particular relevance here, increased amygdala volume has been linked to behavioural inhibition and social anxiety.^68,69^ The amygdala works in concert with pSTS in mediating vigilance for social threat in the external environment.^70^ The pSTS and amygdala are also active during the simulation of social evaluative threat and embarrassment.^71,72^

Here, we used VBM to identify correlations between SRE and/or STE and rGMV across the whole brain. We dissected and quantified the unique and overlapping rGMV correlates of SRE and STE using a combination of raw LODESTARS scores and LODESTARS scores that were orthogonalised (residualised) with respect to one another.

## Methods

### Participants and procedure

A power analysis^73^ indicated a sample size of n = 82 was required to detect a medium sized correlation (r = 0.3, alpha = 0.05, power = 0.8). One hundred right-handed healthy volunteers participated (74 female, 26 male, mean age 24 years, range: 18–54). Participants completed a battery of measures including the LODESTARS, administered using Qualtrics (Provo, UT, http://www.qualtrics.com). Participants attended the imaging centre on a separate occasion for MRI scanning.

### Measuring dispositional social expectancies: The LODESTARS

The LODESTARS is a 10-item inventory examining the extent to which respondents expect to experience social reward (pleasure) and threat (distress) during an imminent vividly imagined social encounter with a group of unfamiliar peers. Participants are asked to imagine that they have joined a new group, club or society and that this evening they will be going to a social event organized by this group/club/society. Participants imagine that this will be the first time they will meet other people who are in the group/club/society. After noting down the name of the group/club/society they have chosen, participants indicate their anticipated and anticipatory cognitions and emotions about the upcoming imagined event, by responding to 10 items on a 5-point Likert scale (see https://osf.io/hq5sg/ for the full measure). Approaching unfamiliar others and establishing initial social connections are core tasks when transitioning into novel social environments (e.g. entering university), and a prerequisite for integrating new people into one’s social network.^1^

Expectancies about social interactions are partly situation-specific;^74^ however, there is a component of them that is influenced by individuals’ temperament and stable working models (schemas) of self and others.^75,76^ The LODESTARS was designed to tap the stable component, by probing participants’ expectancies for interactions with peers (with whom the participant is motivated to interact) in a generic social event context. The scenario described in the LODESTARS is emotionally ambiguous, and thus in line with existing measures in which participants imagine themselves in an emotionally ambiguous (future) scenario.^58,77^ These measures are sensitive to individual differences in affective style.^46,78^ We used an imminent, self-relevant imaginary scenario, since short-term predictions enhance the tendency to rely on episodic, experiential emotional information, relative to personal semantic knowledge (beliefs, traits, etc.).^46,78,79^

Data from more than 1,300 participants demonstrate that the LODESTARS has a two-factor (reward, threat) structure and excellent psychometric properties, including high test-retest reliability.^60,61^ The LODESTARS yields two scores for each participant: a social reward expectancy (SRE) score and a social threat expectancy (STE) score, both of which can range from 1 (low) to 5 (high). The LODESTARS has excellent construct validity and appears to be sensitive in distinguishing different social cognitive-affective processing styles. For example, attachment anxiety is associated with heightened STE, while avoidant attachment is associated with reduced social SRE.^60^ Qualitative data from a community sample confirmed that people find the LODESTARS to be highly naturalistic,^61^ consistent with findings that people devote considerable time in daily life to imagining and evaluating social encounters.^80^

### Image acquisition

T1-weighted anatomical images for each participant were acquired using a 3-T GE HDx MRI scanner at Cardiff University Brain Research Imaging Centre (CUBRIC). The 3-D T1-weighted whole-brain images were acquired using a fast spoiled gradient echo sequence (FSPGR) with 1 × 1 × 1 mm voxel size and between 168 and 182 contiguous slices. Image acquisition parameters were as follows: repetition time (TR) = 7.8 ms echo time (TE) = 2.984 ms; inversion time = 450 ms; flip angle = 15°; data matrix = 256 × 192. These data were usually acquired within one week of the participant completing the LODESTARS (mode = 3 days).

### Image analysis

VBM was performed using SPM12 (Wellcome Trust Centre for Neuroimaging, http://www.fil.ion.ucl.ac.uk/spm/software/spm12) implemented in MATLAB v. R2012b (The MathWorks). First each participant’s structural image was segmented into grey matter (GM), white matter (WM) and cerebrospinal fluid (CSF) using the ‘unified segmentation’ set of algorithms in SPM12. The image segments of interest (the GM segments) were then normalised to MNI space using the diffeomorphic anatomical registration through exponentiated lie-algebra (DARTEL) registration method in SPM12.^81^ The GM images were smoothed using a Gaussian kernel of 8 mm full width at half maximum. An 8mm smoothing kernel is optimal for detecting morpho-metric differences in both large and small neural structures^82^.

### Statistical analysis 1: LODESTARS VBM

We examined correlations between regional grey matter volume (rGMV) and social reward expectancy and social threat expectancies from the LODESTARS. We accounted for the potentially confounding variables of age and gender^83^ by entering them into the general linear models as ‘regressors of no interest’. Participants’ overall brain volumes were also accounted for, by means of proportional scaling in SPM12^84^. A binary MNI brain mask (SPM8 brainmask.nii) was used to restrict the analysed volume to voxels within the brain.

## Model specification

Inference as to whether regional rGMV significantly correlates with one or both regressors of interest requires that *both* LODESTARS-reward and threat scores be included within the same model.^85^

We would not expect reward and threat expectancies to be orthogonal either behaviourally nor necessarily in the brain.^86^ However, it is informative to clarify the effects on rGMV that are uniquely attributable to each of these two regressors. Entering both into a GLM will automatically achieve this: an essential property of the GLM is that only the variability unique to each regressor drives the parameter estimate for it, so that each effect is adjusted for all others.^87,88^ Only assessing the rGMV associations of variance that is unique to threat and to reward carries its own problems however. These are due to the fact that the standard process of GLM parameter estimation removes the effects of shared variability.^87^ When two regressors are highly correlated, their shared variability is large and the unique component for each is correspondingly small. This results in a loss of statistical power. Further, in this case, it is interesting to explore not only the regional rGMV differences uniquely associated with threat or reward expectancies, but also those present when the shared variance is included within the model.

The correlation between LODESTARS-threat and -reward scores in the present study was -.36, *p* = 0.0002 (95% CI = -.56 to -.137), indicating significant shared variance between these two regressors. In order to construct GLMs that incorporate the shared variance component, two new variables were created: LODESTARS-threat orthogonalised with respect to reward (LODESTARS_threat_orth) and LODESTARS-reward orthogonalised with respect to threat (LODESTARS_reward_orth). These variables are the residuals that result from regressing threat on reward and vice versa. By definition, these constitute the portions of each LODESTARS score that are not predicted by the other LODESTARS score.

Using the orthogonalised LODESTARS variables in combination with the ‘raw’ (non-orthogonalised) scores, it was possible to run two GLMs, which between them allowed assessment of individual differences in rGMV uniquely attributable to variance in LODESTARS-threat or reward, as well as rGMV associations present when the shared variance was included but attributed exclusively to threat or reward. I.e., the effects of reward expectancies adjusted for threat and unadjusted for threat, plus the effects of threat adjusted and unadjusted for reward. The two models are specified below. See Figure 1 for a diagrammatic representation of the assignation of (shared) variance that results from orthogonalisation.

**Figure 1:**
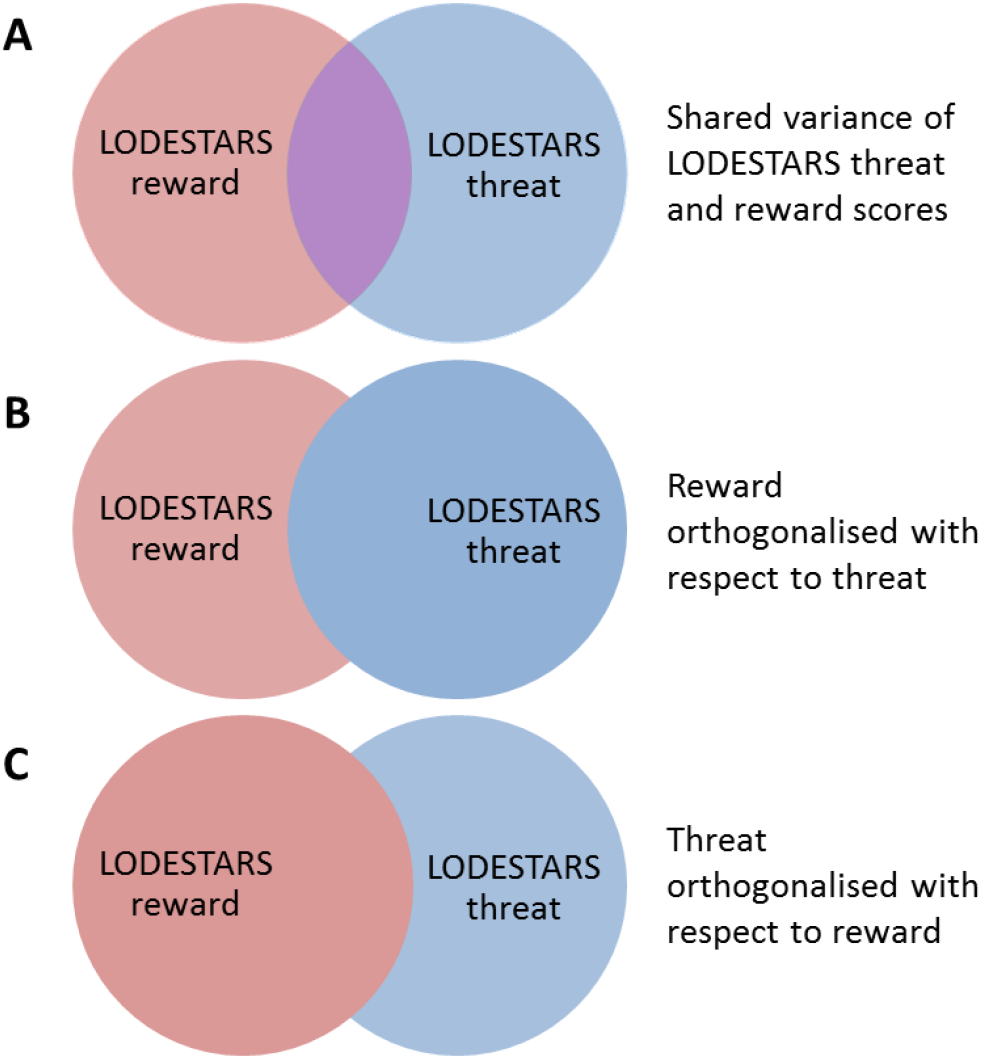
Venn diagrams illustrating how the variability is distributed across the 2 LODESTARS regressors where red is unique to reward, blue is unique to threat and purple is shared. A depicts ‘raw’ LODESTARS-threat and reward scores, which exhibit some overlapping variance. B and C depict the two regression models run, demonstrating the effects of variable orthogonalisation. In **B**, all the shared variance is assigned to LODESTARS-threat while in **C**, all shared variance is assigned to LODESTARS-reward.

Model = Threat orthogonalised with respect to reward. All shared variance assigned to reward.

rGMV = α + b_0_ LODESTARS_reward + b_1_ LODESTARS_threat_orth + b_2_ age + b_3_ gender

Model = Reward orthogonalised with respect to threat. All shared variance assigned to threat.

rGMV = α + b_0_ LODESTARS_reward_orth + b_1_ LODESTARS_threat + b_2_ age + b_3_ gender

## Correction for multiple comparisons

To correct for multiple comparisons across the whole brain, we applied non-stationary cluster extent correction as implemented in the VBM8 toolbox (http://dbm.neuro.uni-jena.de/vbm/) running in SPM12. We used 3DClustSim (AFNI) to calculate the overall expected voxels-per-cluster threshold for our data, for α = 0.05, p ≤ 0.001, based on the brain mask we used (SPM8 brainmask.nii). This gave an expected cluster size of ≥ 86 voxels.

### Statistical analysis 2: Overlap analysis

To test for brain voxels in which rGMV is significantly correlated (positively or negatively) with threat *and* reward expectancies, two further GLMs were applied. These models each contained only one LODESTARS variable as the regressor of interest. The same thresholding was applied as in statistical analysis 1: *p* < 0.005, with an 86-voxel cluster extent threshold.

These models yielded statistical parametric maps (SPMs) of brain regions in which rGMV correlated positively with reward, positively with threat, and negatively with threat. (No clusters survive threshold for negative correlation with reward). These gave rise to two overlap analyses: 1, {reward-positive and threat-negative} and 2, {reward-positive and threat-positive}.

The combinations of SPMs were inspected for overlap by means of masking in SPM12.

### Statistical analysis 3: Structural covariance analyses

To further characterize the network affinities of regions linked with SRE and STE, we examined grey matter structural covariance (SC)^89^ between dmPFC and vmPFC, between vmPFC and amygdala, and between pSTS and amygdala.

We extracted GMVs for the peak voxels of the dmPFC, vmPFC and pSTS clusters that survived cluster-extent correction in the LODESTARS VBM. These voxels were used as seeds in the subsequent analysis.

Our target regions of interest (ROIs) were specified by masks created from the Neuromorphometrics atlas.^90^ Two masks were created: a bilateral amygdala mask and a bilateral vmPFC mask.

We used seed-based SC analyses,^91^ conducted in SPM12, to identify voxels within our target ROIs in which GMV covaried with GMV in the seed voxel. Our analyses identified voxels in which target region GMV covaried positively with seed GMV, and (separately) voxels in which target region GMV covaried negatively with seed GMV. The effects of gender, age, and total brain volume were accounted for in these models. As this was a hypothesis-driven, rather than exploratory analysis, we employed more stringent correction for multiple comparisons than in analyses 1 and 2. Specifically, threshold-free cluster enhancement (TFCE), which controls the family-wise error rate at p < .05.^92^

## Results

The mean LODESTARS-reward score in this sample was 3.7 (from a max. possible score of 5; range = 2.0–4.8); std. dev. = .49) and the mean LODESTARS-threat score was 3.3 (range = 1.0–5.0, std. dev. = .92). Cronbach’s alpha was .65 for LODESTARS-reward and .87 for LODESTARS-threat. There were no significant gender differences in the LODESTARS scores. LODESTARS-reward scores did not correlate with age, however LODESTARS-threat scores decreased with increasing age (*r* = -.30, *p* = .003, 95% CI = -.49 to -.103). This is consistent with findings in a larger sample (n > 1,300).^61^

Both LODESTARS reward and threat scores were significantly higher than the scale midpoint in this sample: for reward, *t* = 14.67, *p* < .001; for threat, *t* = 4.02, *p* < .001. A paired-samples *t*-test indicated that the mean LODESTARS SRE score was significantly higher than mean LODESTARS STE score, *t* = 3.05, *p* = .003, d_av_ = 0.5.

### Statistical analysis 1: LODESTARS VBM results

First, correlations between rGMV and LODESTARS-threat/reward were examined in the SPM T-maps in which shared variance was included. That is, the outputs of the threat orthogonalised with respect to reward model were inspected for correlations between rGMV and LODESTARS-reward scores. The outputs of the reward orthogonalised with respect to threat model were inspected for correlations between rGMV and LODESTARS-threat scores. Details of the clusters that survived non-stationary cluster extent correction are given in Table 1. The extent to which the correlations within each cluster reflect unique variance of threat or reward was then assessed by checking whether the clusters survived cluster-extent correction thresholding for the equivalent contrasts in the opposite model (i.e. reward correlation contrasts in the reward orthogonalised with respect to threat model). These results are reported in the right-most column of Table 1.

**Table 1.**
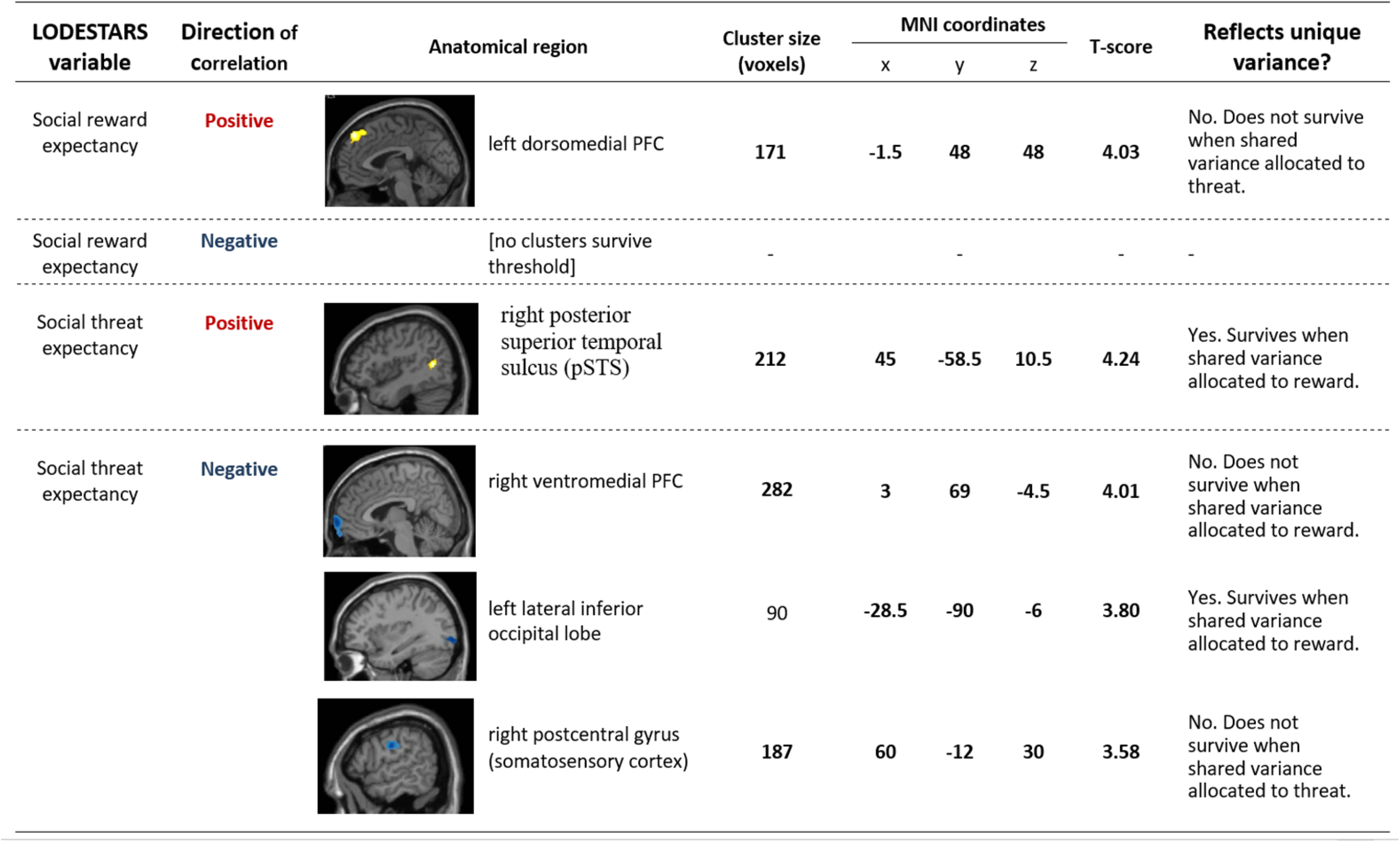
Clusters that survived nonstationary cluster extent correction: shared variance between threat and reward included.

| LODESTARS variable | Direction of correlation | Anatomical region | Cluster size (voxels) | MNI coordinates |  |  | T-score | Reflects unique variance? |
| --- | --- | --- | --- | --- | --- | --- | --- | --- |
|  |  |  |  | x | y | z |  |  |
| Social reward expectancy | Positive | left dorsomedial PFC | 171 | -1.5 | 48 | 48 | 4.03 | No. Does not survive when shared variance allocated to threat. |
| Social reward expectancy | Negative | [no clusters survive threshold] | - | - | - | - | - | - |
| Social threat expectancy | Positive | right posterior superior temporal sulcus (pSTS) | 212 | 45 | -58.5 | 10.5 | 4.24 | Yes. Survives when shared variance allocated to reward. |
| Social threat expectancy | Negative | right ventromedial PFC | 282 | 3 | 69 | -4.5 | 4.01 | No. Does not survive when shared variance allocated to reward. |
|  |  | left lateral inferior occipital lobe | 90 | -28.5 | -90 | -6 | 3.80 | Yes. Survives when shared variance allocated to reward. |
|  |  | right postcentral gyrus (somatosensory cortex) | 187 | 60 | -12 | 30 | 3.58 | No. Does not survive when shared variance allocated to threat. |

A positive correlation between SRE and rGMV was found in a dorsomedial region of left prefrontal cortex (dmPFC, see Figure 2; Figure S1A). This result was significant only in the model in which the shared variance was allocated to reward however; it did not remain significant (at the cluster-size-corrected level) in the model in which the shared variance is allocated to LODESTARS-threat, indicating that this rGMV-expectancy association is partially attributable to shared variance between reward and threat expectancies. No other correlations (positive or negative) of rGMV with LODESTARS-reward survived cluster extent correction.

**Figure 2:**
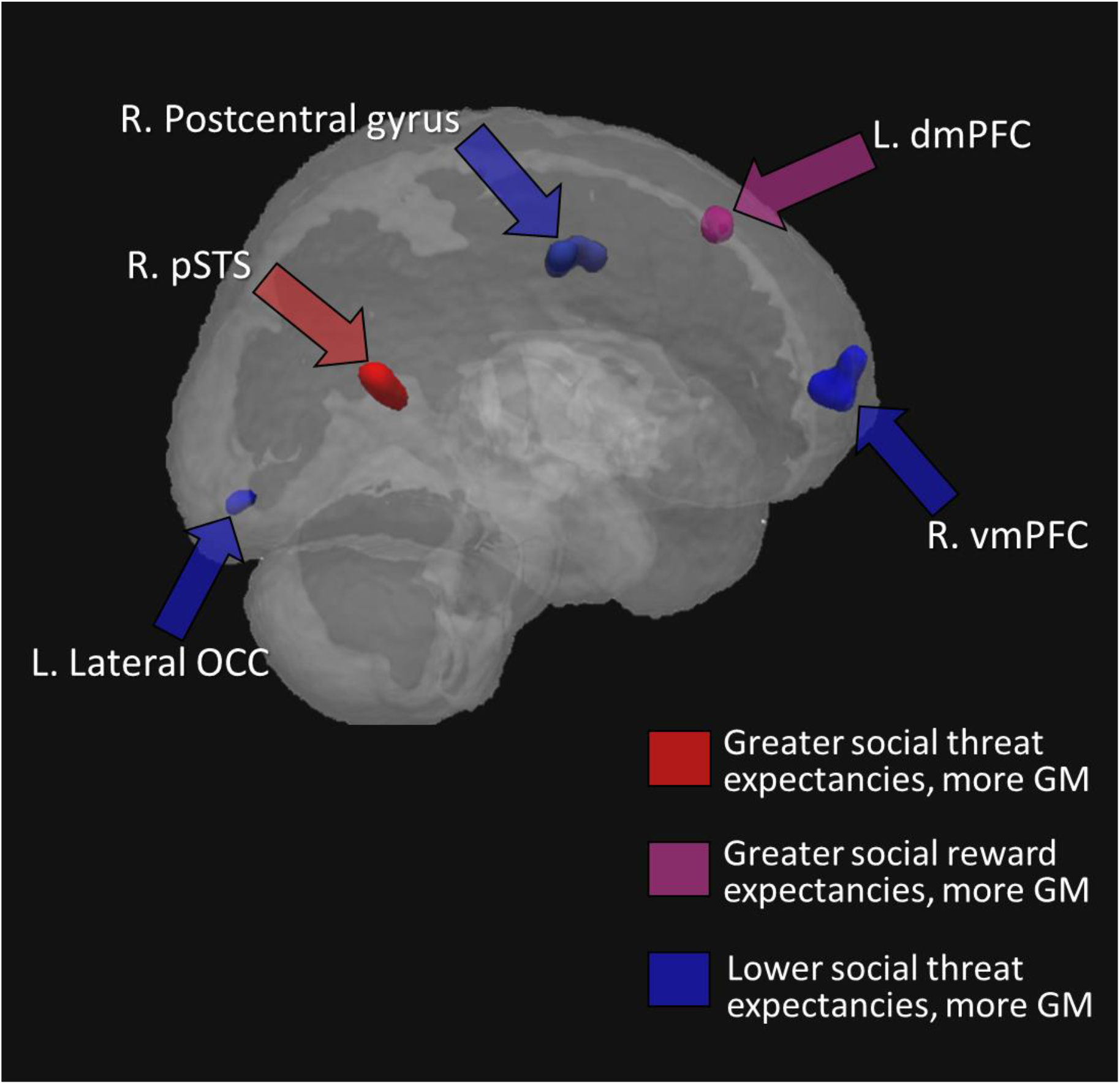
Brain regions in which there were significant associations between self-reported social expectancies and rGMV. For display purposes the clusters are shown at a threshold of p < .001, uncorrected.

Greater rGMV in right posterior superior temporal sulcus (pSTS) was associated with higher STE (Figure 2; Figure S1B), whereas individuals who reported lower expectancies of social threat had greater GM volumes in right ventromedial PFC (vmPFC, see Figure 2; Figure S2A), left lateral occipital lobe (lOCC, see Figure 2; Figure S2B), and right postcentral gyrus (somatosensory cortex, Figure 2; Figure S2C).

The extent and location of each of the clusters that survived non-stationary extent correction are summarised in Figure 2.

### Statistical analysis 2: Brain regions in which rGMV is correlated with both reward and threat expectancies

The results of these overlap analyses are given in Table 3 and Figure 4. The only pairing for which there were overlapping clusters (at p < 0.005, with 86 voxel extent threshold) was {reward-positive and threat-negative}. There was overlap between clusters in the vmPFC (Fig. 4A), in the right lateral inferior temporal gyrus (Fig. 4B) and in right parahippocampal gyrus.

**Table 3.** Overlap of clusters reflecting motivational salience that survived p <0.005, 86-voxel extent threshold.

| Anatomical region | Extent of overlap (voxels) | MNI coordinates |  |  | T-value |
| --- | --- | --- | --- | --- | --- |
|  |  | x | y | z |  |
| right ventromedial prefrontal cortex | 68 | 9 | 48 | -21 | 3.41 |
| right lateral inferior temporal gyrus | 90 | 61.5 | -13.5 | -36 | 3.39 |
| right parahippocampal gyrus | 19 | 27 | -25.5 | -31.5 | 3.04 |

### Statistical analysis 3: Structural covariance analyses

Seed-based SC revealed that pSTS rGMV covaried positively with right amygdala rGMV, while vmPFC rGMV covaried negatively with right amygdala rGMV (Figure 3A). rGMV in the dmPFC seed covaried positively with rGMV in vmPFC (Figure 3B).

**Figure 3:**
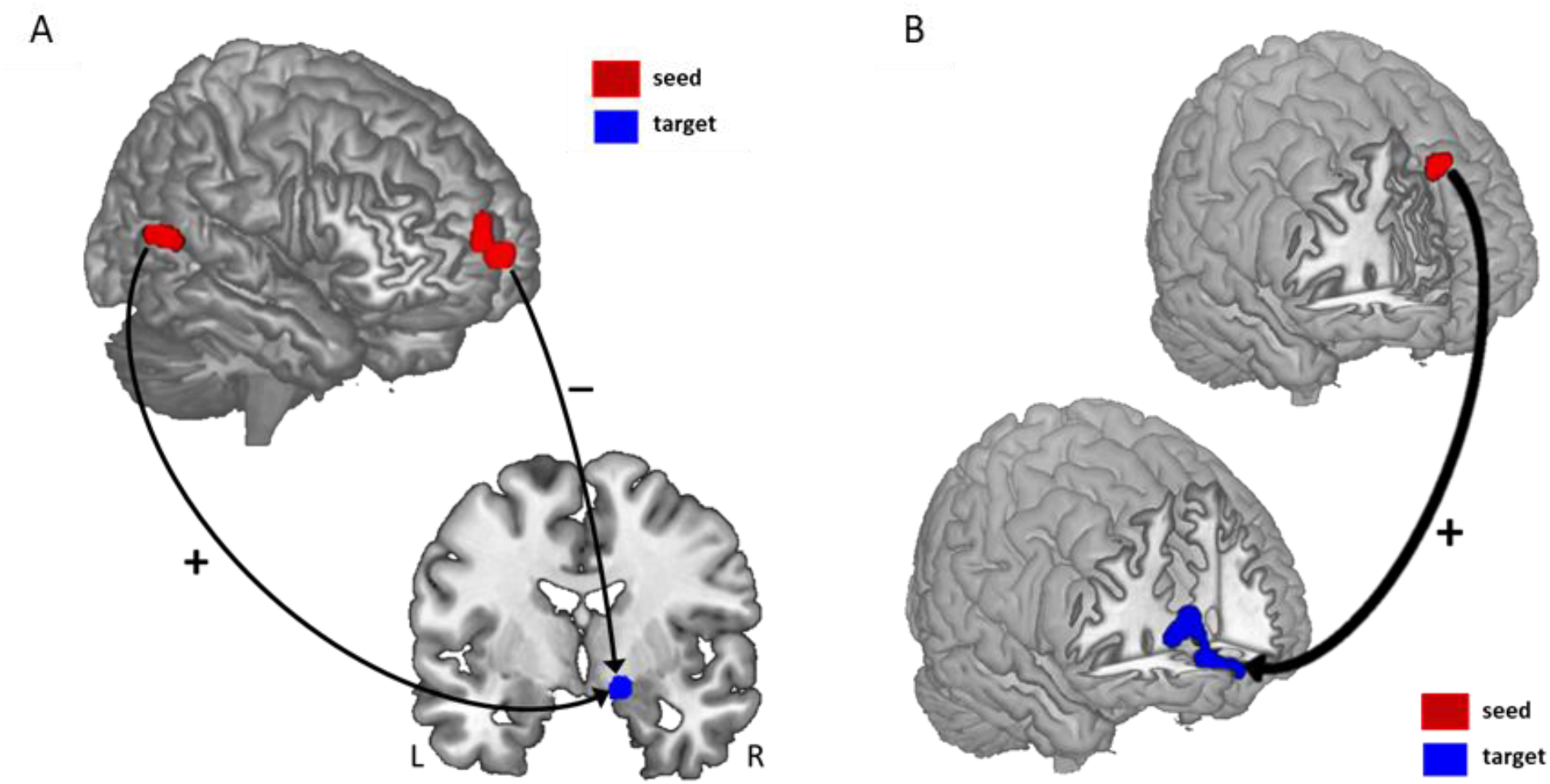
Structural covariance results. **[A]** rGMV in the vmPFC and pSTS seed regions covaried with rGMV in the right amygdala. vmPFC and amygdala rGMV were negatively correlated, while pSTS and amygdala rGMV were positively correlated. **[B]** rGMV in the dmPFC seed region covaried positively with rGMV in the vmPFC.

**Figure 4:**
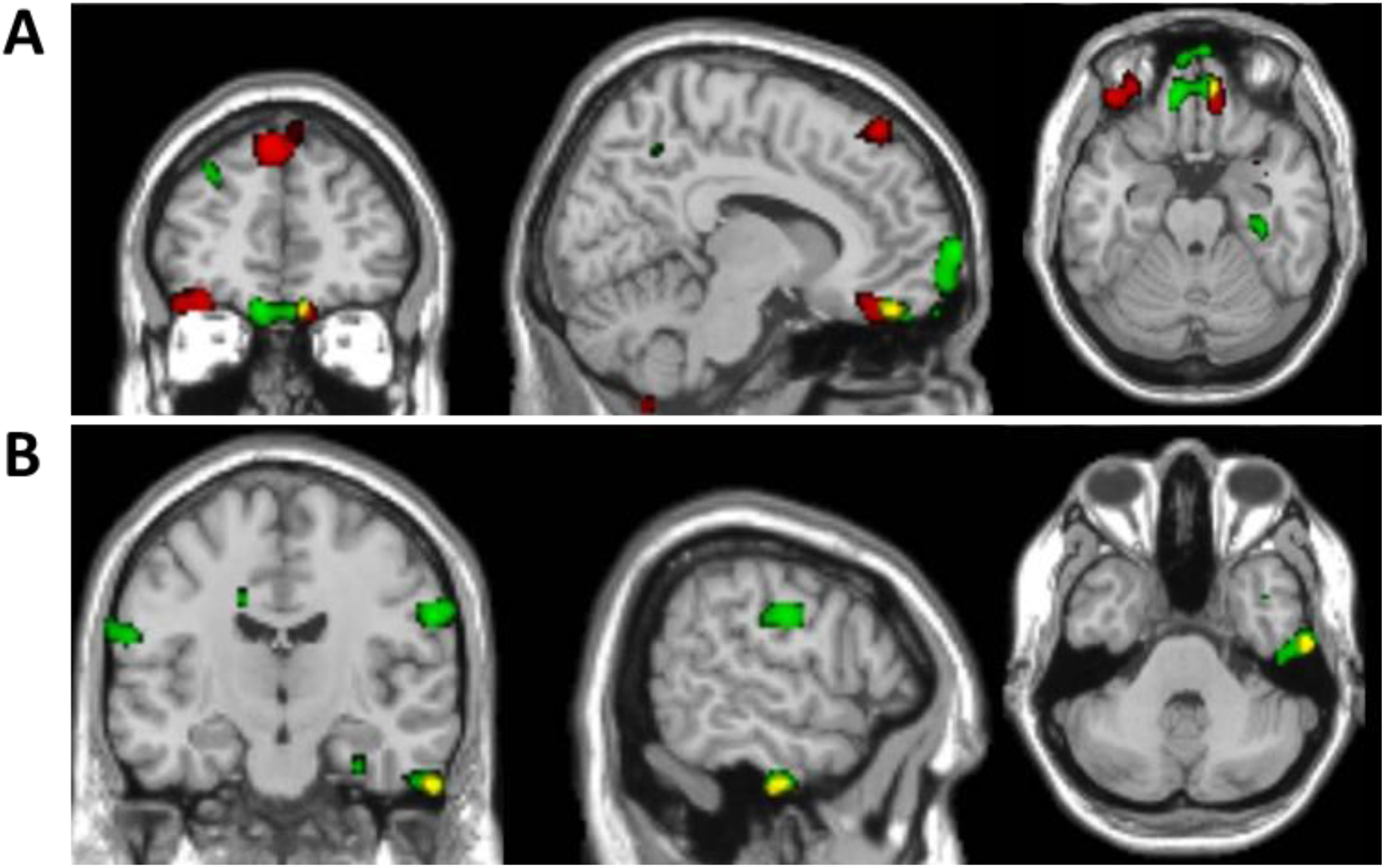
Overlay of regions in which rGMV correlates positively with social reward expectancy and negatively with social threat expectancy. Red = reward_positive; green = threat_negative; yellow = overlap. The SPMs were thresholded at *p* < 0.005 with 10 voxel minimum cluster extent. **A** (upper panel) shows the extent of overlap in right orbitofrontal/ventromedial prefrontal cortex. **B** (lower panel) shows the overlap in right lateral inferior temporal gyrus.

## Discussion

We report a set of focal brain regions in which regional grey matter volume (rGMV) is associated with individual differences in dispositional expectancies of social reward or threat. The results were consistent with previous functional studies revealing that individual differences in future-oriented emotions are underpinned by a network centred on vmPFC.^93^ Further, seed-based structural covariance analyses were consistent with the suggestion that networks anchored in the amygdala support unique dispositions for fostering and maintaining social relationships.^94^

Our novel scenario-based measure generated considerable individual differences in both reward and threat expectancies for the imagined social event. Social reward expectancies (SRE) were significantly higher than social threat expectancies (STE). This finding is robust (n > 1,300^61^) and in line with previous research showing that healthy young adults typically anticipate social acceptance and positive social evaluation from novel interpersonal interactions (e.g.^53,95–97^) as part of a more general optimistic view of their personal future.^46^ This optimism bias is considered to be adaptive,^98,99^ beneficial for physical health and vital for mental health.^100,101^ SRE were largely independent of STE, although the two were modestly inversely correlated (see also ^102^).

Several prominent models posit that two neurobehavioral systems underlie individual differences in affect and motivation.^103,104^ Prospection is at the heart of these models. The appetitive (or approach) system underlies reward pursuit, in part by generating anticipatory and/or anticipated positive emotions. The aversive system underlies anxiety, vigilance, and withdrawal (behavioural inhibition) at the prospect of threat. Our findings align with these models and are broadly consistent with other research showing that social approach and avoidance motives – characterized as the ‘hope for affiliation’ and ‘fear of rejection’ respectively – are distinct dispositions.^15,24^ Further, our work and others’ indicates that positive and negative future-related cognitions are best conceived as separate dimensions of experience, differentially associated with anhedonia and anxiety, respectively.^46,105^

### VBM findings – correlations with social reward expectancies

Previous research shows that anticipated pleasure from imagined social interactions correlates with enhanced vividness of imagined people and places,^106^ and that the spatio-temporal clarity of imagined events is greater for events evoking anticipated positive vs. negative affect.^107,108^ Further, optimism is associated with the tendency to vividly imagine positive events in one’s future (e.g.^109,110^); whereas anhedonia is associated with reduced capacity to simulate detailed future positive events (e.g.^111–114^) as well as reduced accessibility of such images.^46^ Positive episodic expectancy (‘anticipatory savouring’) requires a vivid, contextually detailed mental representation of future reward.

Here, overlap analysis revealed that rGMVs in vmPFC, parahippocampal cortex (PHC) and ventral anterior temporal lobe (vATL) were positively correlated with social reward expectancies *and* negatively correlated with social threat expectancies. These are all regions of a core remembering-imagining network (e.g.^49^). Consistent with our VBM results, these regions are more activated during the simulation of positive, rather than negative, future events.^115^ As part of this network, vmPFC tracks the anticipated positive affective quality of future scenarios,^93,116–119^ consistent with a broader role in subjective valuation of both actually experienced and mental simulated events.^120,121^

vmPFC tracks subjective value as a function of one’s needs and chronic goals^122,123^ and is sensitive to individuals’ optimism bias in their *expectancies* about the hedonic rewards or other benefits that the participant hopes to obtain from such events.^124^ For example, the level of vmPFC activity when imagining positive vs. negative future scenarios is positively correlated with trait optimism.^118^ Of particular relevance to our findings, vmPFC activity to anticipated *social* feedback is enhanced when participants have positive expectancies about social outcomes^96,125,126^ (see also ^115^).

Functionally, vmPFC interacts with PHC and vATL to produce structured positively-valenced mental representations replete with detailed spatiotemporal context and vivid (personal) semantic and sensory details.^93^ Our findings that rGMV not only in vmPFC, but also PHC and ATL, is higher in individuals with higher social reward expectancies is congruent with the behavioural work cited above and further relates to the finding of reduced engagement of these regions during prospection in patients with depression.^127^

Positively biased simulations are partly grounded in biased encoding, consolidation and/or retrieval of autobiographical memories.^128,129^ Speer et al.^130^ found increased dmPFC activity linked to recall of positive autobiographical memories (‘savouring’). Our finding of greater dmPFC rGMV in people with more positive expectancies further corroborates the neural entwining of expectancies and memories.^131^ Our finding of positive structural covariance between dmPFC and vmPFC – which likely reflects long term increased functional connectivity^89^ – may be because the social context inherent in positive mental constructions enhances their value.^126^ It is also possible that the reward value of a simulated event may motivate the degree to which participants engage in mentalizing processes subserved by dmPFC.^132^

rGMV in vmPFC and PHC were also correlated with lower STE. Reduced vividness of positive future thinking is characteristic of anxious individuals, in addition to anxious expectancies about future social interactions.^113,133^ Social anxiety can be regarded as a position along a continuum ranging from a lack of anxiety, to mild shyness and then social anxiety disorder (SAD),^134,135^ so our findings can meaningfully be compared with studies of SAD, which show reduced vmPFC volume.^62,136^

The correlation of vmPFC rGMV with lower STE and greater SRE concurs with the well-established role of vmPFC in emotion regulation. A large-scale neuroimaging meta-analysis of affect regulation across 3 distinct domains (fear extinction, placebo effects, cognitive reappraisal) identified vmPFC activation as the only ‘common neural regulator’ dampening current and anticipated negative affect^66^ (see also ^137^).

These results support the hypothesis that vmPFC plays a ubiquitous role in dampening current and anticipated negative affect.^66^ Our data extend previous work by indicating that the minimisation of STE – and/or the maintenance low threat expectancies – may be implemented in the brain by similar means as the reduction of fear or negative affect in other emotion regulation scenarios.

In healthy adults, successful down-regulation of negative affect is consistently associated not only with increased BOLD activity in the vmPFC, but also with concordant reduction of activity in the amygdala.^66,138–141^ Structural connectivity strength (via white-matter pathways) between vmPFC and amygdala has also been found to be inversely correlated with trait anxiety.^142,143^ Our structural covariance findings add further convergent evidence of the regulatory link between these regions by demonstrating a negative correlation between amygdala rGMV and vmPFC rGMV.

### VBM findings – correlations with social threat expectancies

There were several unique rGMV correlates of individual differences in STE. Heightened anxious (threat) expectancies (fears of potential embarrassment and social rejection) were associated with increased rGMV in right pSTS, alongside decreased rGMV in somatosensory-related cortex (SRC) and lateral occipital cortex (OCC).

Cognitive theories posit that heightened social anxiety results from biased information processing^144^. Alongside regulatory deficits, a processing style marked by hypervigilance and an attentional bias to the social environment for signals of social evaluation is considered a causal and maintaining factor in social anxiety.^145^

Our results are in line with studies suggesting that pSTS serves as an interface between perception of social information and social cognition.^146–148^ pSTS plays a role in analysing socially relevant perceptual information (eye gaze, tone of voice, facial and bodily threat signals), evaluating its implications and orienting attention accordingly, in line with the individual’s present affective state and social goals.^149–151^ pSTS rGMV is increased in SAD and shyness (e.g. ^152,153^), and increased pSTS activity to social perceptual cues (eye gaze etc.) has been consistently demonstrated in individuals who are social inhibited, shy, and socially anxious.^154–159^ Further, resting amygdala–pSTS functional connectivity has been linked to biased social attention and perception in social anxiety.^146,160^ Collectively, this work suggests that chronic hypervigilance for threat may result from, or result in, increased rGMV in right pSTS. Increased expectancies of threat when anticipating future situations may be fundamentally underpinned by these attentional biases.^144,161,162^

Heightened attention to threat may lead to enhanced encoding, elaboration, consolidation and retrieval of negatively biased memories,^163^ resulting in an increased tendency to construct negatively biased expectancies. Further, increased internal attention to threat may maintain attention to negatively constructed future simulations in spontaneous thought, leading to heightened subjective expectancies of their occurrence and increased anticipatory worry.^164^ In turn, this may lead to repercussive effects with increased expectancies further increasing biased attention.^162^

Our findings support cognitive models such as the constructive episodic simulation hypothesis,^165,166^ as we show that the neural structures underpinning attentional biases also underpin prospective ones. Other research has found that pSTS activity is related to remembering and imagining socially threatening situations;^71,72^ and is increased during such simulations in individuals with SAD.^167^

Surprisingly, we *did not* find that amygdala volume directly correlates with individuals’ STE, despite its well-established role in threat processing, including anticipation of social evaluation^168^ and a proposed role in mediating temperamental shyness.^169^ However, we did find positive structural co-variation of pSTS with amygdala, and negative structural covariance of vmPFC and amygdala, consistent with their bidirectional anatomical connectivity.^142,170^

We also found reduced rGMV in left lateral OCC, a region that, together with fusiform gyrus, pSTS and amygdala, forms a face perception network.^171,172^ This may link to fMRI work showing increased pSTS activity to face emotion, but decreased OCC activity (alongside poor face identity recognition) in socially inhibited individuals.^155,173^

Somatosensory-related cortex (SRC) plays a key role in interoception.^174^ Our finding of greater SRC rGMV associated with lower STE thus align closely with findings that individuals with reduced interoceptive sensitivity report significantly greater uncertainty and worry in anticipation of public speaking.^175^ Increased uncertainty in social situations may arise not just because of reduced ability to represent/regulate one’s own interoceptive signals, but also because SRC plays a role in automatic affective empathy via simulation of others’ bodily states. Personal distress (a dysfunctional form of empathy linked with maladaptive emotion regulation and social avoidance^176^) has been shown to be linked to lower rGMV in SRC.^177^

Together, the rGMV correlates of STE we find concur with cognitive models of anxiety, in particular, the combined model,^178^ which contends that socially anxious persons simultaneously exhibit altered processing of internal (distress) cues and external stimuli potentially indicative of negative evaluation.

### Limitations

There are some limitations that should be considered when interpreting our results.

Our study was cross-sectional and so cannot determine whether the relationships between rGMV, SRE and STE arise over time through experience-dependent brain plasticity, or alternatively whether individuals with a specific brain structure are predisposed to acquire different expectancies. Most likely, our findings reflect complex gene-environment interactions over development. In future, training studies such as ^179^ could address this.

The cellular basis of rGMV differences identified by VBM is still poorly understood.^180^ Any tissue property (e.g. cell density, cell size, myelination) that affects relaxation times, and hence voxel images on T1-weighted MRI, will influence VBM measures.

Finally, the generalizability of our results is unknown. We deliberately chose to study a population of university students, because of the ecological relevance of joining new social groups.^1^ Additionally, each participant imagined just one scenario. The scenario was designed to be both sufficiently specific to allow episodic simulation whilst sufficiently generic, such that generalized expectancies (e.g. beliefs) could be tapped. Previous studies (e.g. ^102^), however, suggest a marked degree of consistency across social situations in reward/threat expectancies.

### Conclusions

We found that individual differences in future-oriented thinking in the social domain are reflected in brain macrostructure. In particular, the extent to which individuals hold optimistic vs. pessimistic expectancies for the hedonic outcomes of an imagined social interaction is reflected in rGMV of key regulatory regions, most notably vmPFC. Our findings concur with the idea that vmPFC may integrate various sources of information to conceive the meaning of events for one’s well-being and future prospects.^63^ Our results may reflect a neural embedding of such self-related affective valuation, perhaps accounting for the link between vmPFC macrostructure and adaptive social functioning and well-being.^181,182^

## Supporting information

Supplementary results figures

## Acknowledgements

We thank our participants, CUBRIC core staff for scanning support, and Prof. Yu-Chen Chan for advice on creating the structural covariance figures.

