## Supplementary results figures for "Individual differences in social reward and threat expectancies linked to grey matter volumes in key regions of the social brain"

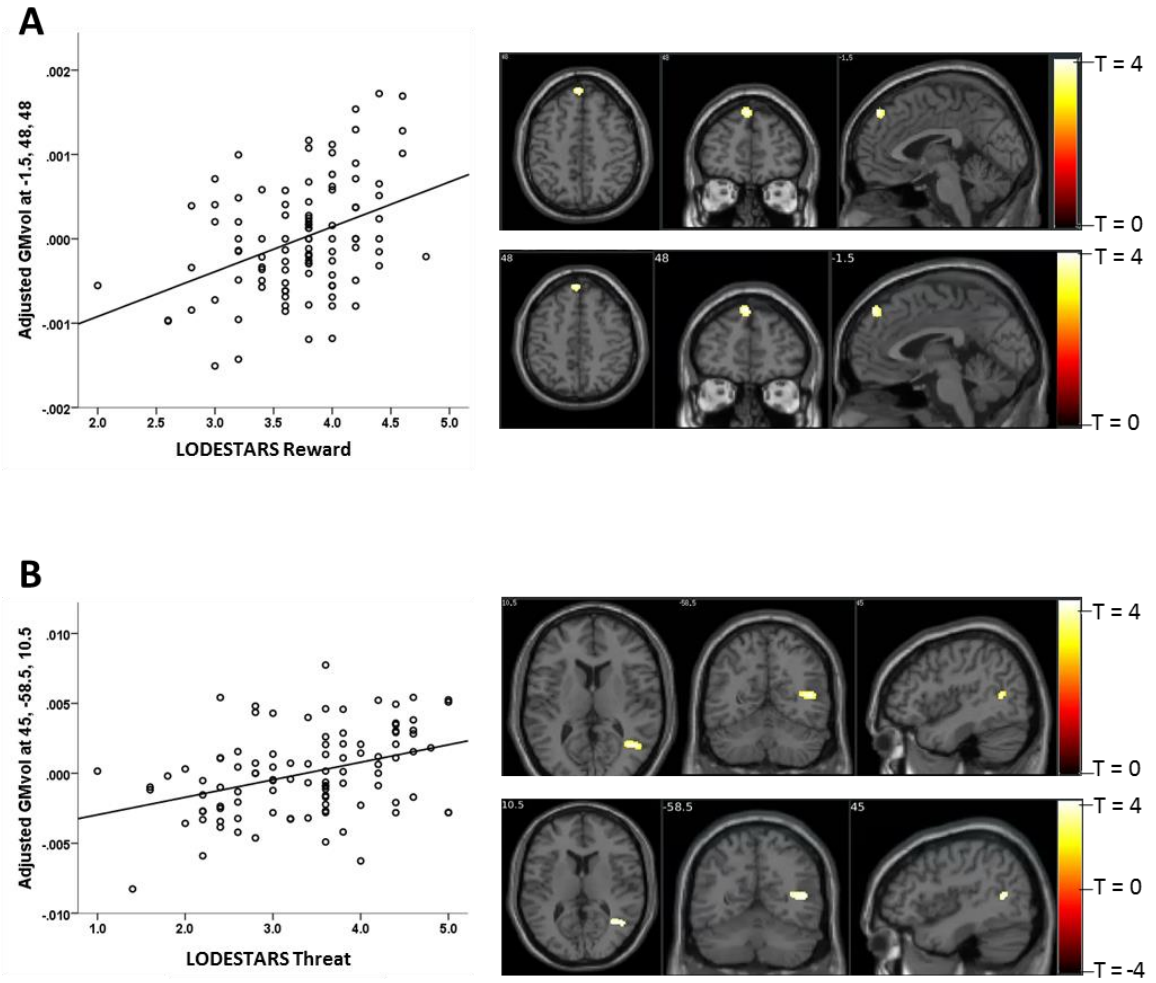


**Figure S1:** Positive correlations between GMvol and LODESTARS scores.

**A.** Greater GMvol in left dorsomedial prefrontal cortex (dmPFC) associated with higher expectations of social reward.

**B.** Greater GMvol in right posterior middle temporal gyrus and superior temporal sulcus (pMTG/STS) associated with higher expectations of social threat.

In both A and B:

Upper panel: cluster-extent corrected. Lower panel: extent of the cluster at $p < 0.001$, unc.

Scatterplot shows the relationship between adjusted GMvol at the peak voxel in the cluster with LODESTARS scores.

Correlation plots are for illustrative purposes only.

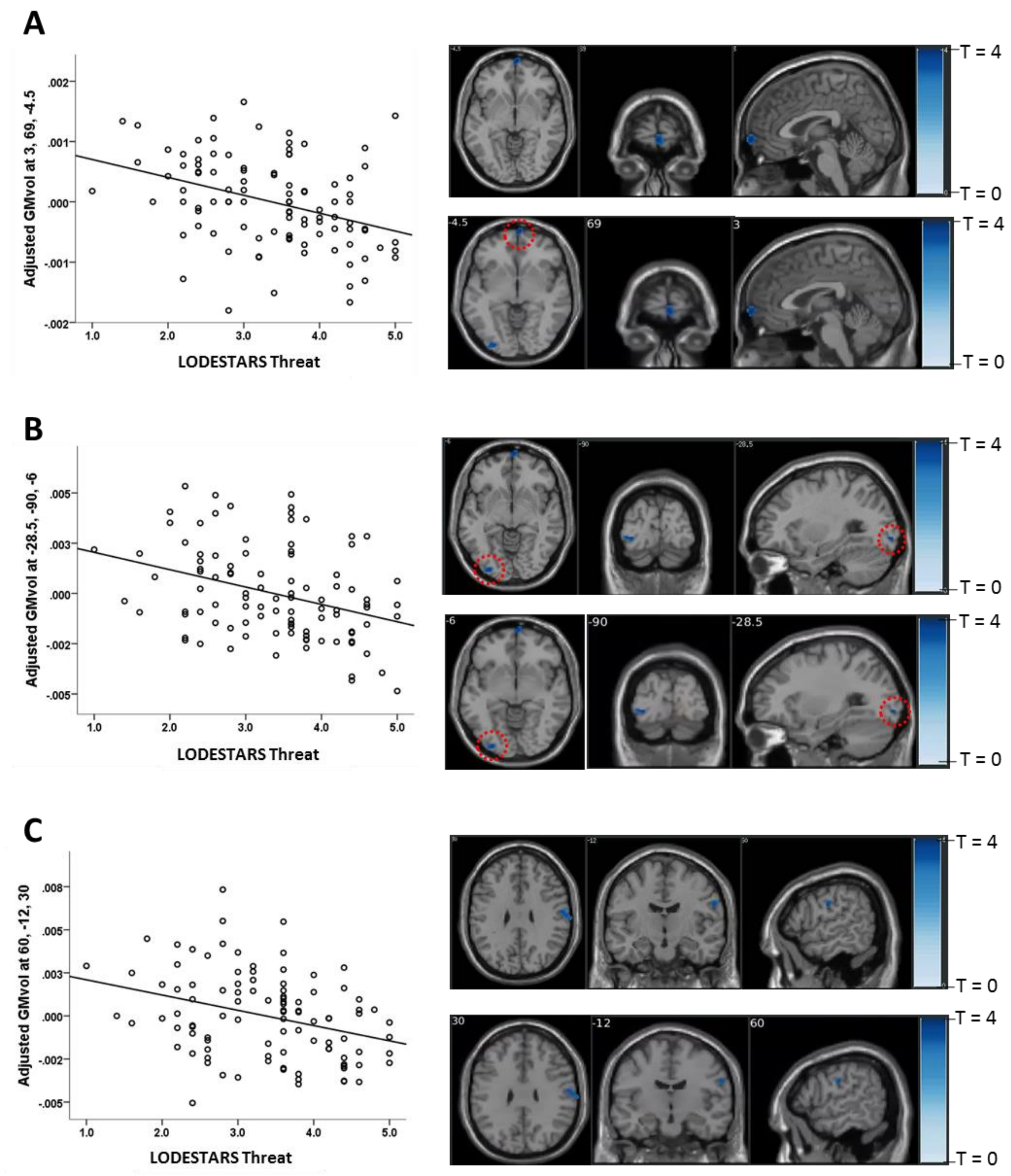


**Figure S2:** Negative correlations between GMvol and LODESTARS scores.

**A.** Greater GMvol in right ventromedial PFC was associated with lower expectations of social threat.

**B.** Greater GMvol in left lateral occipital cortex associated with lower expectations of social threat.

**C.** Greater GMvol in right postcentral gyrus (somatosensory cortex) associated with lower expectations of social threat.

In A, B and C:

Upper panel: cluster-extent corrected. Lower panel: extent of the cluster at $p < 0.001$, unc. Scatterplot shows the relationship between adjusted GMvol at the peak voxel in the cluster with LODESTARS-threat scores.Correlation plots are for illustrative purposes only.
